# Presynaptic GABA_A_ receptors control integration of nicotinic input onto dopaminergic axons in the striatum

**DOI:** 10.1101/2024.06.25.600616

**Authors:** Samuel G. Brill-Weil, Paul F. Kramer, Anthony Yanez, Faye H. Clever, Renshu Zhang, Zayd M. Khaliq

## Abstract

Axons of dopaminergic neurons express gamma-aminobutyric acid type-A receptors (GABA_A_Rs) and nicotinic acetylcholine receptors (nAChRs) which are both independently positioned to shape striatal dopamine release. Using electrophysiology and calcium imaging, we investigated how interactions between GABA_A_Rs and nAChRs influence dopaminergic axon excitability. Direct axonal recordings showed that benzodiazepine application suppresses subthreshold axonal input from cholinergic interneurons (CINs). In imaging experiments, we used the first temporal derivative of presynaptic calcium signals to distinguish between direct- and nAChR-evoked activity in dopaminergic axons. We found that GABA_A_R antagonism with gabazine selectively enhanced nAChR-evoked axonal signals. Acetylcholine release was unchanged in gabazine suggesting that GABA_A_Rs located on dopaminergic axons, but not CINs, mediated this enhancement. Unexpectedly, we found that a widely used GABA_A_R antagonist, picrotoxin, inhibits axonal nAChRs and should be used cautiously for striatal circuit analysis. Overall, we demonstrate that GABA_A_Rs on dopaminergic axons regulate integration of nicotinic input to shape presynaptic excitability.

## INTRODUCTION

Midbrain dopaminergic (DA) neurons form highly branched axons that densely innervate the striatum (Matsuda et al., 2009; Aransay et al., 2015). DA release from striatal axons has been implicated in motor control and reward processing across species (Schultz et al., 1983, 1997; Cohen et al., 2012; Dodson et al., 2016; Howe & Dombeck, 2016; da Silva et al., 2018; Markowitz et al., 2023; Azcorra et al., 2023). Several studies have demonstrated that various neurotransmitters are poised to locally modulate DA release via axonal receptors (Kennedy et al., 1992; Chen et al., 2001; H. Zhang & Sulzer, 2003; Shin et al., 2015; Sulzer et al., 2016, Roberts et al., 2022). Among these receptors, ionotropic and metabotropic GABA receptors have been shown to influence DA release through multiple mechanisms including via a striatal GABAergic tone and GABA co-release from DA axons themselves (Gruen et al., 1992; Smolders et al., 1995; Pitman et al., 2014; Lopes et al., 2019; Kramer et al., 2020; Holly et al., 2021; Patel et al., 2024). Past work using direct axon recordings demonstrated that GABA_A_ receptors are present on the axons of DA neurons and can inhibit striatal DA release by decreasing axonal excitability and hindering action potential (AP) propagation (Kramer et al., 2020). However, how GABA_A_ receptor signals interact with signaling from other axonal receptors remains an open question.

Nicotinic acetylcholine (ACh) receptors (nAChRs) on the axons of DA neurons are activated by release from striatal cholinergic interneurons (CINs) which provide another form of local control over striatal DA release (H. Zhang & Sulzer, 2004; Rice & Cragg, 2004; Threlfell et al., 2012; Cachope et al., 2012; Mamaligas et al., 2016; C. Liu et al., 2022; Kramer et al., 2022). Synchronous activation of CINs causes robust DA release in *ex vivo* preparations and recent work has shown that activation of single CINs is sufficient to evoke small amounts of DA release (Threlfell et al., 2012; C. Liu et al., 2022; Matityahu et al., 2023). Data from direct axonal recordings has shown that DA axons exhibit spontaneous, nAChR-mediated depolarizations as a result of transmission from nearby CINs (Kramer et al., 2022). The subthreshold depolarizations can summate in the axon to evoke axonal APs and drive DA release (Zhou et al., 2001; Yorgason et al., 2017; C. Liu et al., 2022; Kramer et al., 2022). Thus, the ability to regulate the amplitude of nicotinic depolarizations would provide a means of controlling the excitability of DA axons and subsequent neurotransmitter release.

Previous work has shown that the GABA_A_ positive allosteric modulator (PAM) diazepam reduces evoked DA release moderately in the presence of nAChR antagonists but inhibits DA release to a much larger extent when nicotinic transmission is left intact (Kramer et al., 2020). Although the mechanism is unclear, this observation raises the possibility that interactions between GABA_A_ and nicotinic receptors may shape DA release. In particular, it is well-known that GABA_A_ receptors on the soma and dendrites can dampen neuronal excitability and limit the integration of excitatory inputs (Staley & Mody, 1992; Chance et al., 2002; Mitchell & Silver, 2003; Farrant & Kaila, 2007; Lipkin & Bender, 2023). Whether axonal GABA_A_ receptors play a similar role to shape nAChR-mediated input onto DA axons has not been examined.

Here, we tested the influence of axonal GABA_A_ receptors in controlling integration of nicotinic input within DA axons. Using direct axonal recordings, we show that GABA_A_ receptor signaling suppresses subthreshold nicotinic input in individual DA axons. We make use of genetically encoded calcium (Ca^2+^) indicators (GECIs) with fast on-kinetics to record electrically-evoked, multi-component Ca^2+^ influx in DA axons. We show that the first derivative of these signals allows for unambiguous quantification of two temporally-distinct components; one arising from direct DA axon stimulation and the other mediated by nAChR activation. Application of the GABA_A_ receptor antagonist gabazine enhances the nAChR-mediated component of Ca^2+^ influx with only modest effects on the direct component Ca^2+^ influx and no effect on ACh release. Lastly, we show that the commonly used GABA_A_ receptor antagonist picrotoxin inhibits nAChRs at concentrations that are typically used in physiology experiments. Altogether, we demonstrate that axonal GABA_A_ receptors impose a brake on the integration of nicotinic input in DA axons to shape presynaptic calcium signals and axonal excitability.

## RESULTS

### Potentiation of GABA_A_ receptor activation with diazepam suppresses nicotinic receptor-mediated EPSPs in individual DA axons

Striatal DA axons receive tonic GABA_A_ receptor input and phasic nAChR input via ACh release from CINs (Lopes et al., 2019; Kramer et al., 2020; Kramer et al., 2022; **Fig. 1A**). Previous experiments conducted in the presence of nAChR antagonists to isolate the effects of GABA_A_ receptors showed that evoked DA release is reduced by bath application of the GABA_A_ receptor PAM diazepam (Kramer et al., 2020). Interestingly, diazepam results in stronger inhibition of DA release when nicotinic transmission is left intact but the mechanisms that underlie this phenomenon are not understood.

**Figure 1:**
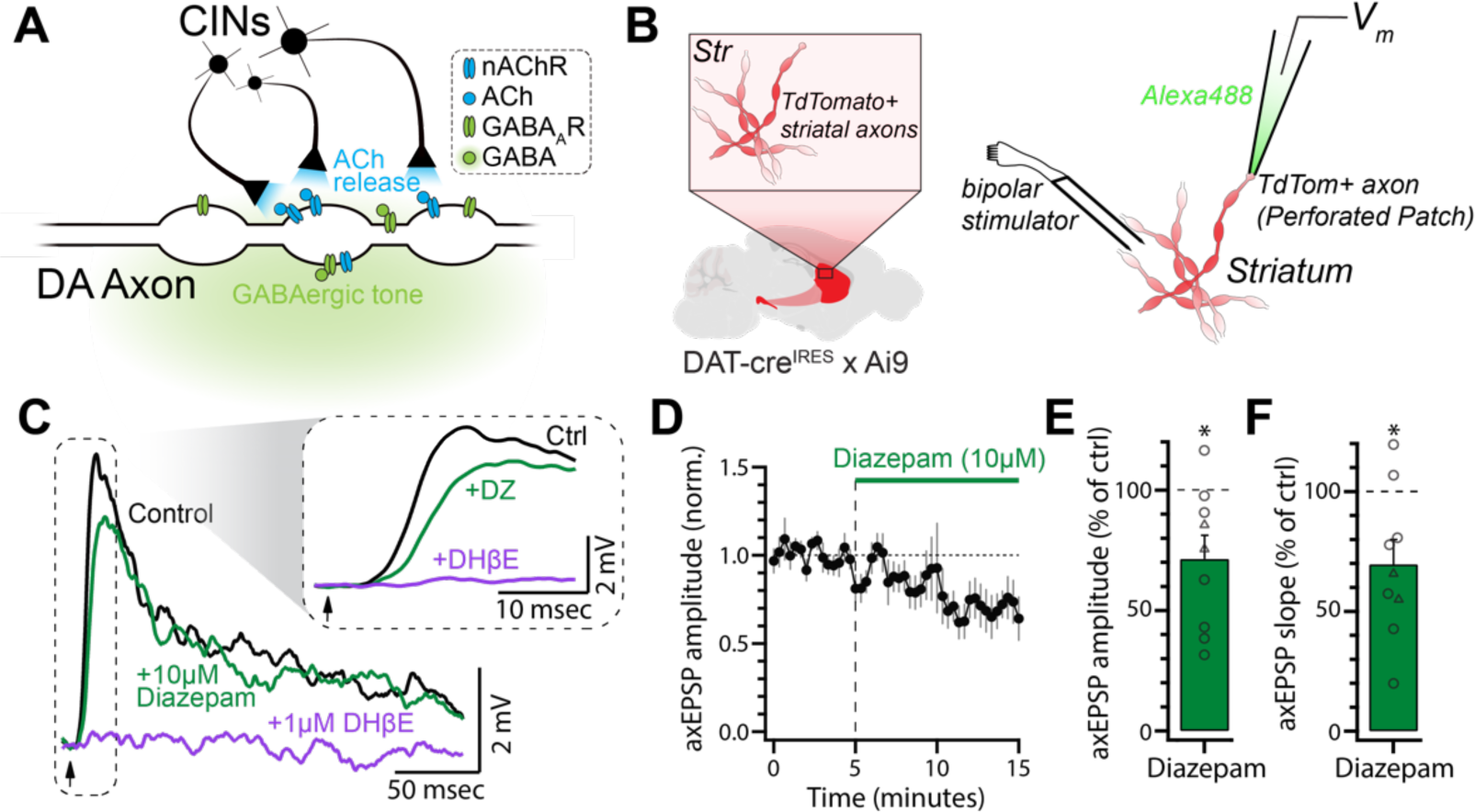
Diazepam suppresses subthreshold nicotinic input onto DA axons. **(A)** Illustration of the simplified striatal environment that DA axons are subjected to. Namely, a GABAergic tone that acts upon GABA_A_ receptors and ACh release from CINs that acts on nAChRs. **(B)** Schematic of recording set-up. Low-intensity electrical stimulation was applied in the vicinity of an axon recorded in perforated patch configuration. **(C)** Average electrically-evoked axEPSPs from a representative axon during control conditions, after diazepam (10 μM) wash-in, and after DHβE (1 μM) wash-in. **(D)** Normalized amplitude of evoked axEPSPs during diazepam wash-in (n = 9). **(E)** Quantification of data in (C) (n = 9). **(F)** The effect of diazepam on the rising slope of axEPSPs. Slope was measured between 30% and 70% of the peak amplitude (n = 9). In a subset of recordings (circles in E and F; triangles represent axons recorded in normal ACSF), nAChRs were partially blocked with a sub-saturating concentration of hexamethonium chloride (200 μM) to ensure that the amplitude of axEPSPs remained subthreshold.

We examined the effects of diazepam on nAChR-evoked electrical signaling in DA axons. To do so, we performed direct recordings from DA axons in the dorsomedial striatum (DMS) in horizontal slices (**Fig. 1B**). Perforated patch recordings from striatal DA axons were used to examine electrically-evoked CIN transmission and limit rundown of the nicotinic input (Wu et al., 2004; Kramer et al., 2022). Consistent with our past work, we found that low-intensity electrical stimulation resulted in nAChR-mediated axonal EPSPs (axEPSPs) that were blocked by dihydro-β-erythroidine hydrobromide (DHβE, 1 μM; Kramer et al., 2022; **Fig. 1C**). Following wash-in of diazepam (10 µM), we observed that the amplitude of evoked axEPSPs were reduced by ∼28% (71.40 ± 9.77% of control, p = 0.0190, n = 9; **Fig. 1C-D**). In addition, the rising slope of the axEPSPs was significantly slower in diazepam when compared to control (69.64 ± 10.30% of control, p = 0.0185, n = 9; **Fig. 1F**). Importantly, these effects were observed without exogenously applied GABA_A_ receptor agonists, which is consistent with the presence of a striatal GABAergic tone as previously suggested (Kramer et al., 2020). Together, these data suggest that the amplitude and kinetics of nicotinic input onto individual striatal DA axons are controlled by tonically activated axonal GABA_A_ receptors.

### Electrically-evoked Ca^2+^ signals from DA axons contain temporally- and pharmacologically-distinct components

To further investigate interactions between GABA_A_ receptors and nAChRs, we next tested the excitability of DA axons at the population level using GECI signals. Specifically, we expressed the high-affinity jGCaMP8s in DA neurons via viral injections in DAT-Cre mice (Y. Zhang et al., 2023; **Fig. 2A**). To record spike-evoked Ca^2+^ signals in DA axons, we used single pulses of local, near-maximal electrical stimulation with bipolar electrodes placed in striatal slices and measured fluorescence with a photodiode (**Fig. 2A-B**). We found that application of DHβE (1 μM) to block nAChRs dramatically reduced the peak jGCaMP8s signal by ∼88% compared to the original value (DHβE: 12.16 ± 2.79% of control, p < 0.0001; n = 6; **Fig. 2C**). Subsequent addition of tetrodotoxin (TTX; 500 nM) completely abolished the remaining evoked Ca^2+^ signal (TTX: 0.79 ± 0.30% of control, p < 0.0001; n = 6; **Fig. 2C**). These data show that evoked Ca^2+^ signals represent axonal APs generated both by direct stimulation of DA axons as well as those generated by axonal nAChR activation (**Fig. 2D**).

**Figure 2:**
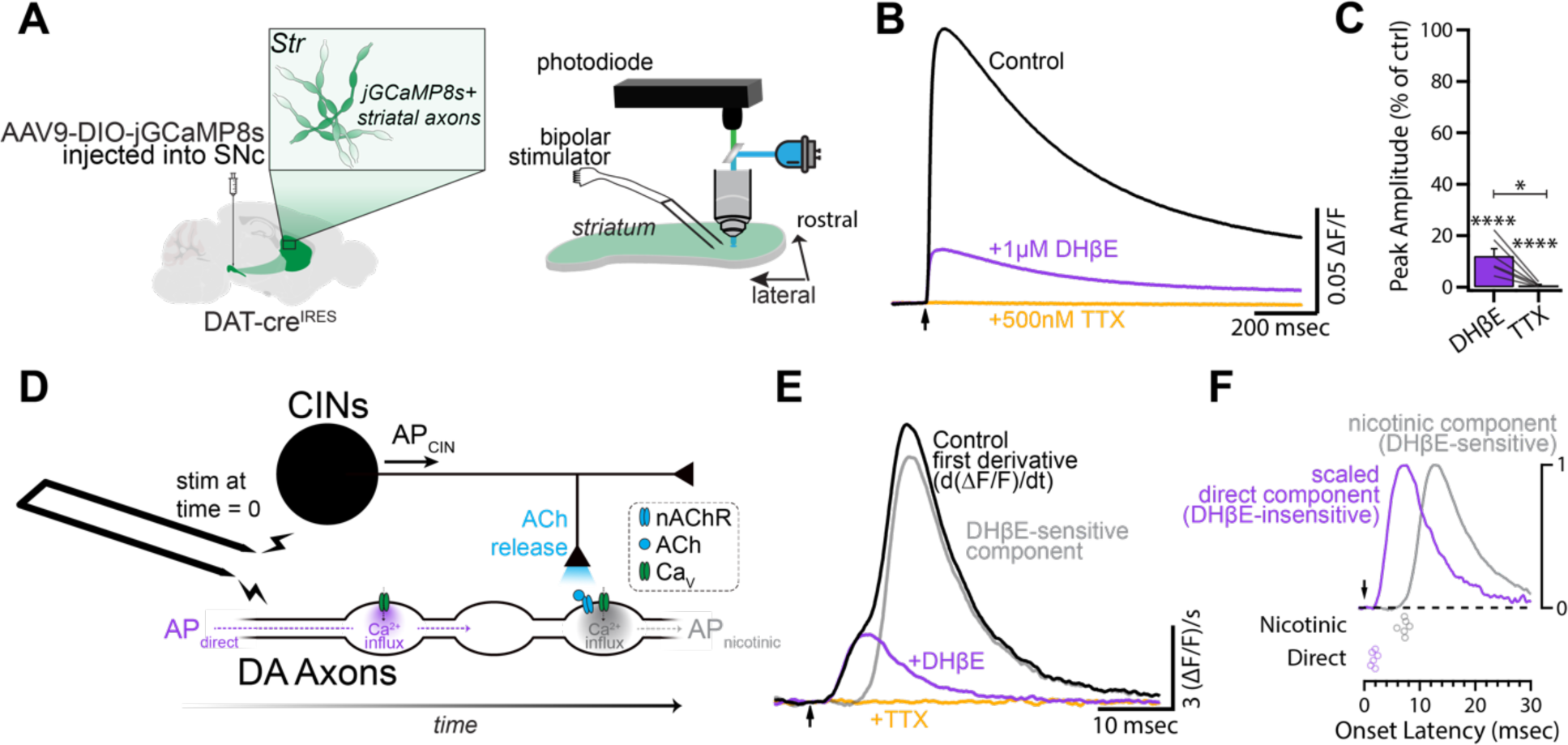
jGCaMP8s as a readout of multi-component DA axon activation. **(A)** Schematic of jGCaMP8s expression strategy and experimental set-up. Horizontal slices containing the dorsomedial striatum were imaged with a photodiode while stimulated with a bipolar electrode 200 μm away and out of the field of view of the photodiode. **(B)** Fluorescent signals following a single electrical stimulus. Each trace represents an average of each drug condition in a representative experiment. **(C)** Normalized amplitude in DHβE (1 μM) and TTX (500 nM) compared to control (n = 6). **(D)** Schematic of CIN-DA axon circuitry and time course of APs following suprathreshold electrical stimulation. **(E)** First derivative of traces in (B). Grey trace is the subtraction of the DHβE trace from the control trace. **(F)** Top: Normalized nicotinic and direct components from (E). Bottom: Onset latencies for the two components (n = 6).

To distinguish between nAChR-mediated and directly-mediated Ca^2+^ influx during jGCaMP8s transients in our photometric data, we calculated the first temporal derivative of the fluorescence signals, as previously done to estimate presynaptic Ca^2+^ currents in cerebellar parallel fibers (Sabatini & Regehr, 1996, 1997, 1998). The first derivative of the evoked DA axon jGCaMP8s fluorescence revealed two components. The first ‘direct component’ occurred immediately following electrical stimulation and was unchanged following blockade of nAChRs (**Fig. 2E, Fig. S1**; see methods). This direct component was TTX-sensitive and had an onset latency of 1.74 msec (SEM: ±0.18 msec, n = 6; **Fig. 2E, F**). This component likely corresponds to the direct activation of DA fibers. The second, DHβE-sensitive ‘nicotinic component’ occurred later in the signal and had an onset latency of 7.12 msec (SEM: ±0.30 msec, n = 6; **Fig. 2F**). This latency measurement is in agreement with previous work that has shown that the latency of CIN transmission onto DA axons is around 7 msec (Wang et al., 2014b; Kramer et al., 2022). The similarity of the waveforms of the two components is consistent with the notion that both reflect temporally-distinct APs in DA axons (**Fig. 2F**). Using these methods in subsequent experiments, therefore, enabled us to measure presynaptic Ca^2+^ influx resulting from both directly-evoked APs and nAChR-evoked APs.

### GABA_A_ receptor antagonism enhances evoked nicotinic signals in dopaminergic axons

Using the derivative to distinguish between direct and nAChR-evoked DA axon activity, we next sought to examine how GABA_A_ receptor activation interacts with evoked nicotinic transmission in a population of DA axons. One possibility is that axonal GABA_A_ receptors could be regulating nAChR input to DA axons under control conditions. To test this, we bath applied the selective GABA_A_ receptor antagonist SR-95531 (gabazine; 10 μM) following a baseline period. Gabazine application resulted in a significant increase of the electrically-evoked jGCaMP8s signal compared to control (115.59 ± 5.21% of control, p = 0.0202, n = 8; **Fig. 3A-C**). Taking the first derivative revealed that, under these conditions, the potentiation of the jGCaMP8s signal was primarily driven by a significant increase in the magnitude of the nicotinic component (121.69 ± 7.82% of control, p = 0.0276, n = 8; **Fig. 3D-F**). In contrast, the direct component exhibited a small but non-significant increase in magnitude (107.67± 5.31% of control, p = 0.1922, n = 8; **Fig. 3E, F**). This result is in agreement with our previous findings that tonic GABA has only a modest effect on locally evoked DA release in the presence of nAChR antagonists (Kramer et al., 2020). Additionally, treatment with gabazine resulted in a shorter latency to peak for the nicotinic component, without affecting onset latency of the direct component (Direct: -0.16 ± 0.11 msec, p = 0.1928; Nic: -0.59 ± 0.24 msec, p = 0.0431; n = 8; **Fig. 3G, Fig. S2**).

**Figure 3:**
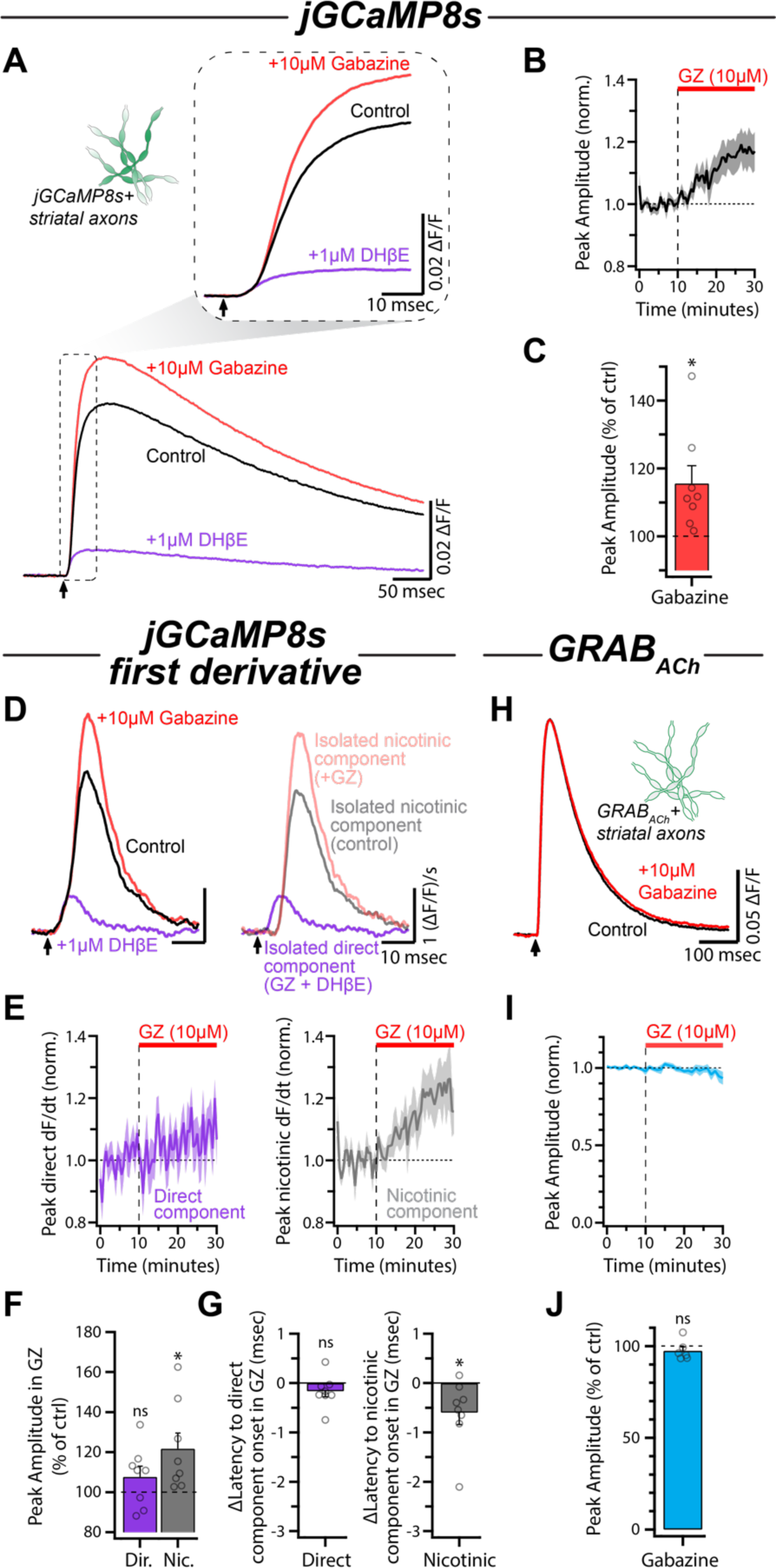
GABA_A_R antagonism potentiates the nicotinic component without affecting ACh release. **(A)** Average jGCaMP8s signals for each condition in a representative experiment. Inset shows the same traces on an expanded timescale. **(B)** Time course of the peak amplitude during gabazine (10 μM) wash-in (n = 8). **(C)** Quantification of the peak amplitude of the jGCaMP8s signals in gabazine compared to control. **(D)** Left: First derivative of traces in (A). Right: Direct component and nicotinic components overlaid. **(E)** Time course of normalized direct component and nicotinic component amplitude during wash-in of gabazine. **(F)** Normalized amplitudes of both components in gabazine compared to control. **(G)** Latency of the two components in gabazine compared to control (see also Fig. S2). **(H)** Average evoked GRAB_ACh_ signals during control conditions and following wash-in of gabazine for a representative experiment. **(I)** Time course of the normalized amplitude of GRAB_ACh_ signals during gabazine wash-in (n = 6). **(J)** Normalized amplitude of GRAB_ACh_ signals in gabazine compared to control.

The changes to the nicotinic component in gabazine could be explained simply by disinhibition of CINs which results in an increase of ACh release. To test this hypothesis, we examined the effect of gabazine directly on ACh release from CINs using the fluorescent ACh sensor GRAB_ACh3.0_ (abbreviated as GRAB_ACh_) virally expressed on DA axons (**Fig. 3H**; Jing et al., 2020). Under the same conditions as the experiments described above, we found that blockade of GABA_A_ receptors with gabazine had no effect on the evoked GRAB_ACh_ signal (amplitude: 97.46 ± 2.19% of control, p = 0.2990, n = 6; **Fig. 3I, J**). These data suggest that GABA_A_ receptors located on the axons of DA neurons suppress the integration of nicotinic input in a manner that is independent from altering ACh release from CINs.

### Picrotoxin robustly inhibits nicotinic receptors on dopaminergic axons

Previous work in the field has used a combination of gabazine and picrotoxin to probe the effects of GABA_A_ receptor activation. To test the individual contributions of the two drugs, we applied picrotoxin to slices already bathed in gabazine. In contrast to gabazine alone which enhanced the jGCaMP8s signal, we were surprised to see that subsequent application of picrotoxin (100 μM) resulted in a robust decrease in the evoked jGCaMP8s signal (62.50 ± 3.85% of control, p < 0.0001, n = 8; **Fig. 4A-C**). Comparison of the nicotinic and direct components of the jGCaMP8s derivative signal revealed that the effects of picrotoxin were isolated to the nicotinic component (Direct: 106.30 ± 3.55% of control, p = 0.3534; Nic: 45.76 ± 4.48% of control, p < 0.0001; n = 8; **Fig. 4D-F**). Additionally, this depression of evoked nicotinic input was dose dependent, with an IC_50_ of 107.7 μM (95% confidence interval: 83.94 μM to 144.30 μM; **Fig. 4G**). To examine if these effects resulted from a reduction in presynaptic ACh release, we imaged GRAB_ACh_ in slices already bathed in gabazine (**Fig. 4H**). Addition of picrotoxin to the bath solution did not influence the amplitude of the evoked GRAB_ACh_ signals (amplitude: 99.65 ± 4.09% of control, p = 0.9365, n = 6; **Fig. 4I-J**), indicating that ACh release was not affected by picrotoxin. Together, these results indicate that the effects of picrotoxin on DA axon Ca^2+^ influx are likely due to direct nAChR inhibition.

**Figure 4:**
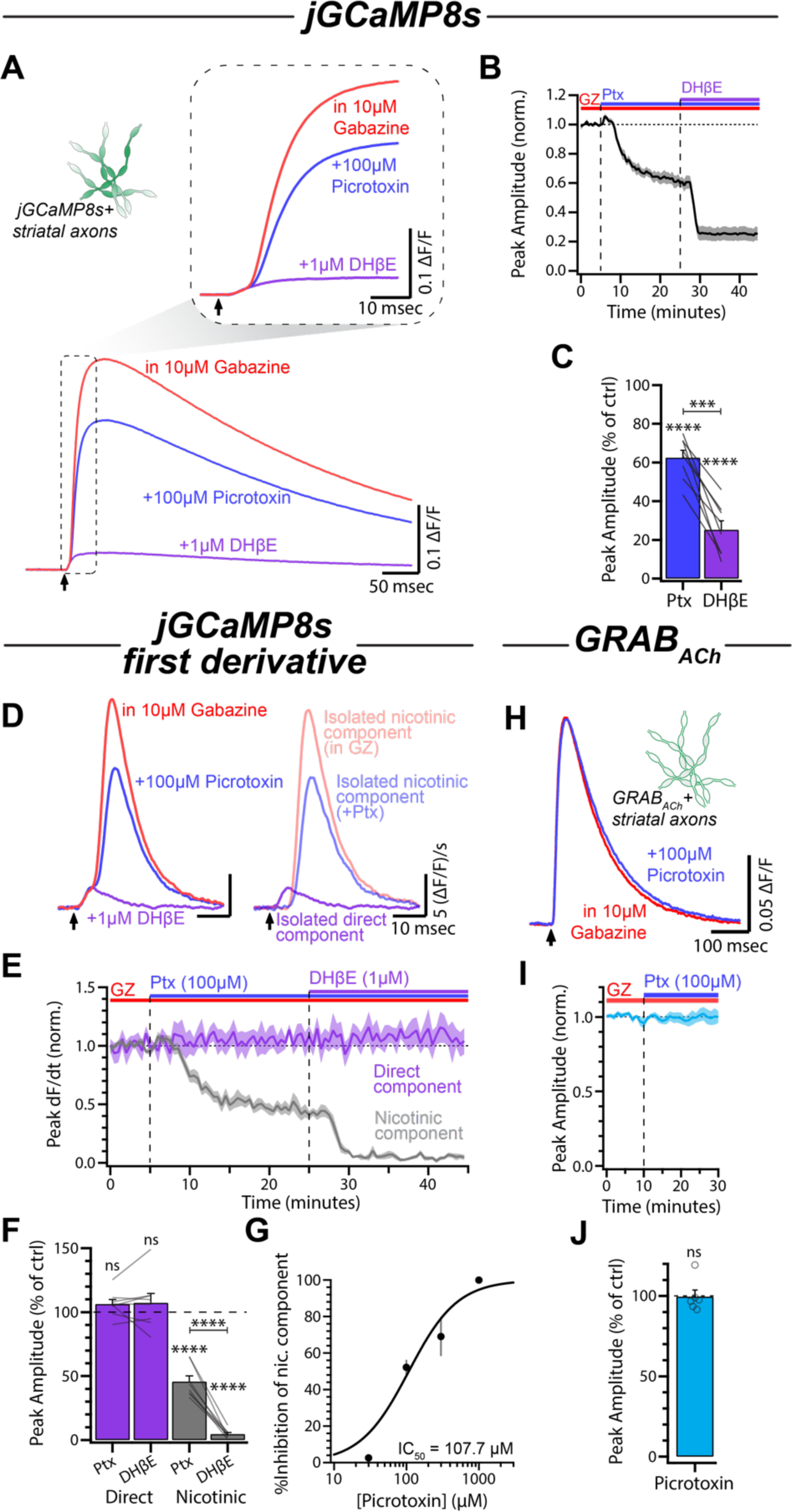
Picrotoxin blocks nAChRs in a dose-dependent manner. **(A)** Average jGCaMP8s signals for each condition in a representative experiment. Inset shows the same traces on an expanded timescale. **(B)** Time course of the peak amplitude during picrotoxin (100 μM) and DHβE (1 μM) wash-in (n = 8). **(C)** Quantification of the peak amplitude of the jGCaMP8s signals in picrotoxin and DHβE compared to control (gabazine, 10 μM). **(D)** Left: First derivative of traces in (A). Right: Direct component and nicotinic components overlaid. **(E)** Time course of normalized direct component and nicotinic component amplitude during wash-in of picrotoxin and DHβE. **(F)** Normalized amplitudes of both components in picrotoxin and DHβE compared to control. **(G)** Dose-response curve of the nicotinic component to increasing picrotoxin concentration (30μM: n = 3; 100 μM: n = 11; 300 μM: n = 5; 1000 μM: n = 4). **(H)** Average evoked GRAB_ACh_ signals in gabazine and following wash-in of picrotoxin for a representative experiment. **(I)** Time course of the normalized amplitude of GRAB_ACh_ signals during picrotoxin wash-in (n = 6). **(J)** Normalized amplitude of GRAB_ACh_ signals in picrotoxin compared to control.

To further rule out circuit interactions as the mechanism for the effects of picrotoxin, we directly tested the effects of picrotoxin on axonal nAChRs using whole-cell axonal recordings from DA axons in the medial forebrain bundle (MFB; **Fig. 5A**). To activate nAChRs, we puff-applied ACh (300 μM) using pressure ejections from a pipette positioned ∼50 μm from the axonal recording site (**Fig. 5A**). ACh puffs produced time-locked depolarizations with amplitudes of ∼10mV (10.51 ± 0.59 mV, n = 6; **Fig. 5B**). Application of picrotoxin (100 μM) reliably reduced the amplitude of the ACh-evoked depolarization by ∼50% (Ptx: 5.27 ± 0.49 mV, p = 0.0018; **Fig. 5B-D**), consistent with direct effects on nAChRs. Subsequent application of DHβE further reduced the depolarizations to amplitudes indistinguishable from baseline fluctuations in membrane potential (DHβE: 1.49 ± 0.23 mV, p = 0.0008; n = 6; **Fig. 5B-D**). Together, these results demonstrate that picrotoxin directly inhibits nAChRs on the axons of DA neurons.

**Figure 5:**
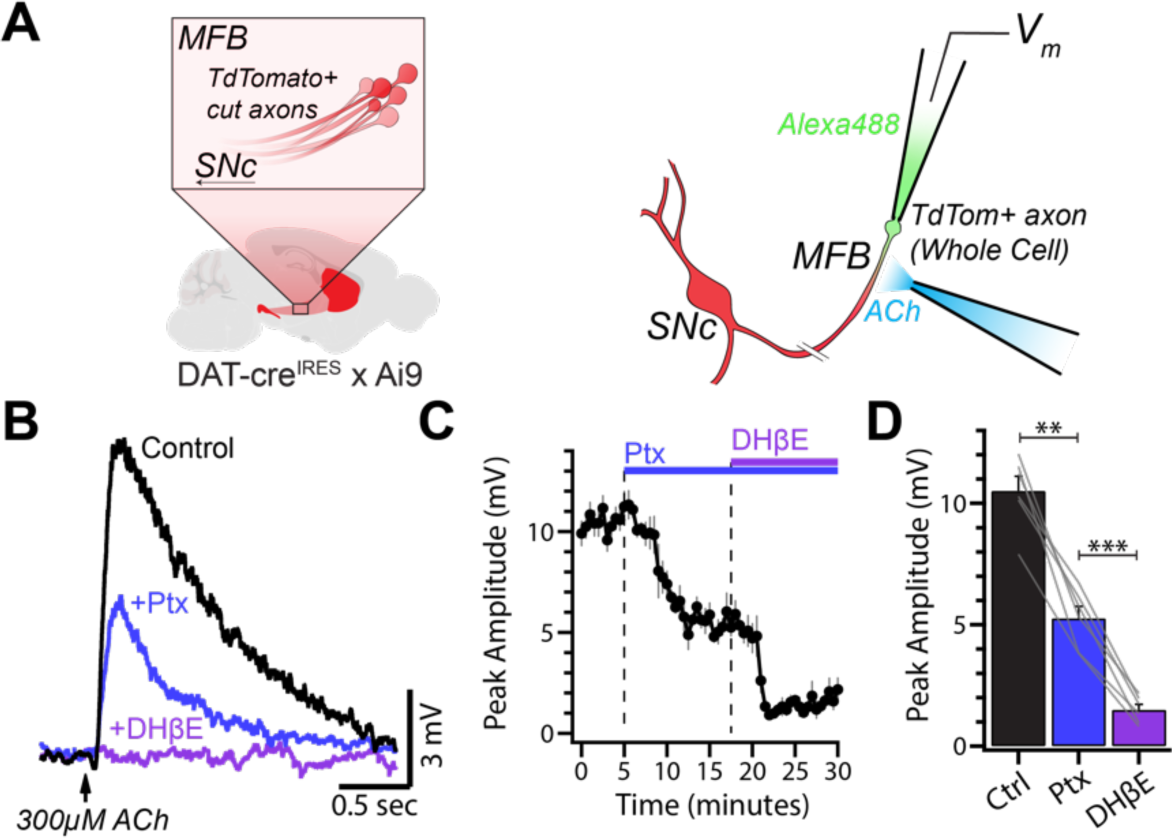
Picrotoxin decreases ACh-evoked depolarizations in the main axon of DA neurons. **(A)** Schematic of recording configuration. ACh (300 μM) was applied from a puff pipette positioned ∼50 μm away from the recording site. **(B)** Average ACh-evoked depolarizations in the main axon of a DA neuron. **(C)** Time course of depolarization amplitude during wash-in of picrotoxin (100 μM) followed by DHϕ3E (1 μM). **(D)** Amplitude of ACh-evoked depolarization in each condition (n = 6).

### Picrotoxin preferentially inhibits non-α6 subunit-containing nicotinic receptors on DA axons

We next tested whether picrotoxin inhibition of nAChRs occurred in a subunit-specific manner. DA axons express both α4 and α6 subunit-containing nAChRs (Zoli et al., 2002). The majority of large-amplitude spontaneous axEPSPs in DA axons are mediated by α6 subunit-containing nAChRs and antagonists for these receptors diminish CIN-evoked DA release (Exley et al., 2008; Wang et al., 2014a; Kramer et al., 2022). Conotoxin-P1A (P1A; 300 nM) was used to selectively block α6 subunit-containing nAChRs (Dowell et al., 2003; McIntosh et al., 2004). Consistent with previous literature examining striatal dop[amine release, P1A application resulted in a decrease of evoked calcium transients in DA axons (30.17 ± 5.48% of control, p < 0.0013, n = 5; **Fig. 6A-C**; Exley et al., 2008; Wang et al., 2014a; Kramer et al., 2022). Specifically, the nicotinic component was reduced by ∼85% while the direct component was not impacted (Nic: 14.79 ± 3.29% of control, p < 0.0001; Direct: 99.23 ± 1.74% of control, p > 0.9999; n = 5; **Fig. 6D-F**). Subsequent application of picrotoxin further diminished the nicotinic component to the same level as DHβE (Ptx: 2.67 ± 0.77% of control, DHβE: 1.34 ± 0.22% of control, p = 0.6835, n = 5; **Fig. 6E, F**). This result is suggestive of a preferential effect on α4 subunit-containing nAChRs.

**Figure 6:**
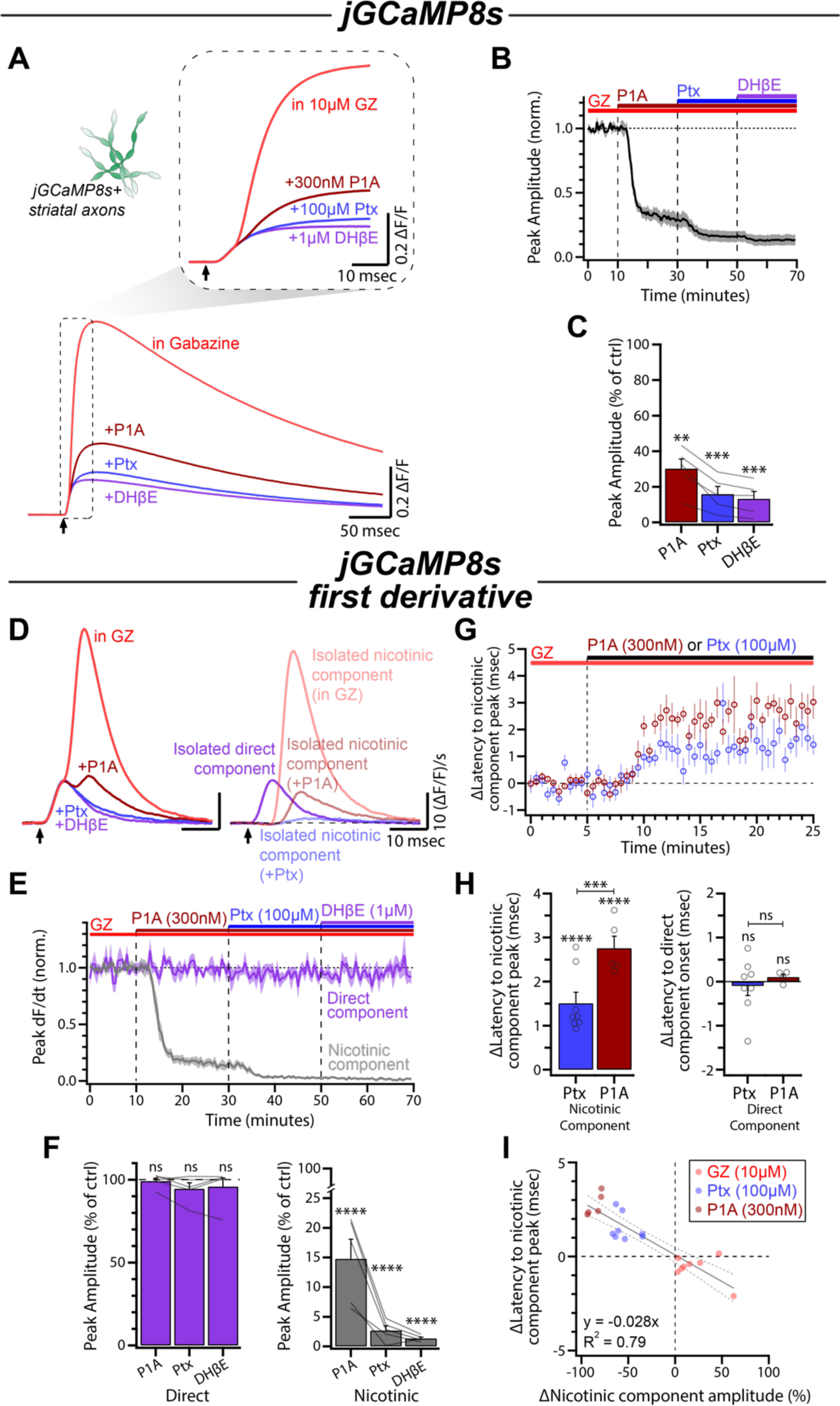
Picrotoxin antagonism of axonal nAChRs is not occluded by conotoxin P1A. **(A)** Average jGCaMP8s signals for each condition in a representative experiment. Inset shows the same traces on an expanded timescale. **(B)** Time course of the peak amplitude during conotoxin-P1A (300 nM), picrotoxin (100 μM), and DHβE (1 μM) wash-in (n = 5). **(C)** Quantification of the peak amplitude of the jGCaMP8s signals in P1A, picrotoxin, and DHβE compared to control (gabazine, 10 μM). **(D)** Left: First derivative of traces in (A). Right: Direct component and nicotinic components overlaid. **(E)** Time course of normalized direct component and nicotinic component amplitude during wash-in of P1A, picrotoxin, and DHβE. **(F)** Normalized amplitudes of both components in P1A, picrotoxin, and DHβE compared to control. **(G)** Time course of the time-to-peak of the nicotinic component following application of either picrotoxin (n = 8) or P1A (n = 5). All slices were bathed in ACSF containing 10 μM gabazine for the duration of the experiment. **(H)** Left: Quantification of data shown in (G). Right: No changes were observed in the onset latency of the direct component in the same experiment (Ptx: n = 8; P1A: n = 4; see also Fig. S3). **(I)** Correlation of effect on nicotinic component amplitude and nicotinic component latency for gabazine, picrotoxin, and P1A (GZ: n = 8; Ptx: n = 8; P1A: n = 5).

Previous work observed both a decrease in axEPSP amplitude and a slowing of axEPSP rise time in P1A (Kramer et al., 2022). Consistent with those results, we observed longer a latency to peak of the nicotinic component upon application of P1A (+2.76 ± 0.27 msec, p < 0.0001, n = 5; **Fig. 6G, H**). Interestingly, picrotoxin (100 μM) had a similar but smaller effect (+1.50 ± 0.25 msec, p < 0.0001, n = 8; **Fig. 6G, H**). There was no observed shift in the latency to the onset of the direct component following either picrotoxin or P1A treatment, suggesting that the effects on timing were limited to the nicotinic component (Ptx: -0.09 ± 0.22 msec, p > 0.9999, n = 8; P1A: +0.10 ± 0.07 msec, p > 0.9999, n = 4; **Fig. 6H, Fig. S3**). These increases in latency following nAChR blockade with either picrotoxin or P1A are opposite of what we observed following GABA_A_ receptor antagonism with gabazine (**Fig. 3G**). In fact, changes in latency correlate strongly with the effects on amplitude by the three drugs, suggesting that perturbations to either nAChR or GABA_A_ receptor activation can alter both the magnitude and the timing of the nicotinic-evoked Ca^2+^ influx in DA axons (R^2^ = 0.79; **Fig. 6I**). Together, these data show that picrotoxin antagonism of nAChRs behaves similarly to other partial nAChR antagonism, and that the drug may be moderately selective for non α6 subunit-containing nAChRs.

## DISCUSSION

In this study, we investigated GABAergic control over DA axon physiology and employ a method to optically study population-level axonal excitability with GECIs. Specifically, we show that axonal GABA_A_ receptors interact with axonal nAChR signaling to gate nicotinic influence over DA axons. Using direct axonal recordings, we demonstrate the broad-spectrum benzodiazepine, diazepam, dampens subthreshold nicotinic input onto DA axons. Our photometry results indicate that antagonism of GABA_A_ receptors potentiates nAChR-mediated input onto the population of DA axons without affecting ACh release in the striatum. Our data suggest that these effects are not mediated by circuit mechanisms but instead result from integration of the two neurotransmitter signals within the DA axons themselves. Additionally, we provide evidence that picrotoxin exerts off-target effects and directly inhibits nAChRs on DA axons at concentrations often used in slice electrophysiology experiments. Lastly, we compared nAChR antagonism with picrotoxin to that of P1A and show that significant picrotoxin-induced inhibition of axonal Ca^2+^ persists after blockade of α6 subunit-containing nAChRs. Together, these results add to our growing understanding of local control over DA axon physiology and suggest that these axons continuously integrate information from the striatal environment.

### Using the derivative of jGCaMP8s signals as a proxy for presynaptic Ca^2+^ influx and axonal excitability

Here, we use the first derivative of signals from a GECI with a fast on-rate as a proxy for presynaptic Ca^2+^ currents (I ^pre^) arising from both directly-evoked APs and nAChR-mediated APs (Sabatini & Regehr, 1998; Y. Zhang et al., 2023; Y. Zhang & Looger, 2023). Previous work has demonstrated that the first derivative of the fluorescent transient from low-affinity Ca^2+^-sensitive fluorophores can give a quantitative prediction for I ^pre^ (Sabatini & Regehr, 1996, 1997, 1998). Our results demonstrate that comparable analyses can be applied to GECIs to allow for relative measurements of changes to multi-component presynaptic Ca^2+^ influx, similar to analyses that have been applied to amperometric recordings of DA release (Wang et al., 2014a; Wang et al., 2014b). One major advantage of this method is the genetic control it provides compared to the use of bulk-loaded calcium indicators, something that is particularly useful in circuitry without a stereotyped architecture, such as the striatum.

It is important to note that Ca^2+^ can enter striatal DA axons from a number of sources including through internal stores, through nAChRs, and through voltage-gated Ca^2+^ channels following subthreshold depolarizations or following APs (Decker & Dani, 1990; Vernino et al., 1992). As with all measurements made using Ca^2+^ sensors, the jGCaMP8s transients recorded here cannot differentiate between these different sources of Ca^2+^. That said, recent work has shown that AP-mediated Ca^2+^ is likely the primary source of presynaptic Ca^2+^ in DA axons (C. Liu et al., 2022). An additional consideration is that, due to the spatially averaged nature of photometry, it is difficult to determine how many individual axons are contributing to the bulk signals. Increases in the magnitude of the jGCaMP8s signal, as seen following application of gabazine, could therefore be due to a larger amount of presynaptic Ca^2+^ influx in the same number of axons, recruitment of additional axons that were previously not firing APs, or a combination of the two. This ambiguity is intrinsic to all photometry methods and concretely addressing it in the future will require optical or electrical measurements from individual axons.

Our data suggest that evoked GECI fluorescence can be used as a proxy for some features of axonal physiology, such as receptor control of presynaptic calcium. While GECIs are largely regarded as slow proxies for neuronal activity, here we show that the high-affinity jGCaMP8s enables distinction between millisecond-scale activity patterns and is readily able to report changes in the kinetics and amplitude of presynaptic Ca^2+^ influx. Applying this analysis to other neuronal populations could be a useful way to probe axonal excitability and examine the effects of axonal receptors on the physiology of those presynaptic compartments.

### Off-target effects of picrotoxin on DA axon nAChRs

Nicotinic receptors belong to the cys-loop family of receptors along with GABA_A_ receptors, glycine receptors, and type-3 serotonin receptors (Sine & Engel, 2006; Miller & Smart, 2010). While homology between cys-loop receptors and the broad effects of some antagonists, such as bicuculline, have been well-described, the effects of picrotoxin are less clear (Matsubayashi et al., 1998; Rothlin et al., 1999; Demuro et al., 2001). Specifically, contradictory effects of picrotoxin on nAChRs have been reported (Q. Y. Liu et al., 1994; Erkkila et al., 2004; Juárez et al., 2014; Pita-Almenar et al., 2014). Here, using both jGCaMP8s photometry and direct axonal recordings, we show that nicotinic input to DA axons is robustly inhibited by picrotoxin.

One possible explanation for variable reports of picrotoxin antagonism of nAChRs is subunit composition of the receptors. In the striatum, DA axon terminals primarily express α4β2 and α6β2 nAChRs (Picciotto et al., 1998; Klink et al., 2001; Zoli et al., 2002). Subunit specificity has been reported for some nicotinic antagonists, and picrotoxin is likely not an exception (Harvey et al., 1996; Zoli et al., 2002). Due to the sensitivity of the nicotinic component to picrotoxin, our results suggest that picrotoxin is effective at inhibiting axonal nAChRs that, based on previous studies, likely contain the β2 subunit (Picciotto et al., 1998; Zoli et al., 2002). Additionally, the effects of picrotoxin were not occluded by antagonism of α6 subunit-containing nAChRs by initial application of P1A. In fact, picrotoxin exerted stronger effects following the application of P1A – blocking ∼85% of the remaining nicotinic component in the presence of P1A compared to ∼55% under control conditions. These data suggest that picrotoxin may be especially effective at blocking the remaining α4 subunit-containing nAChRs. However, due to the non-linear nature of these measurements, confirmation with more direct experiments testing this hypothesis is necessary.

Overall, we observed a strong reduction of nicotinic input to DA axons when picrotoxin was applied at 100 μM, a concentration regularly used to block GABA_A_ receptors in *ex vivo* brain slice experiments. Our results indicate that previous studies using picrotoxin in the striatum as a putative selective GABA_A_ antagonist may be difficult to interpret. That said, when used under conditions where nicotinic transmission is already blocked by a selective antagonist, picrotoxin can yield unambiguous results regarding the block of GABA_A_ receptors (Lopes et al., 2019; Kramer et al., 2020). In general, our data suggest that the use of a more selective GABA_A_ receptor antagonist, such as gabazine, is preferable for studies in the striatum or other circuitry that involves both GABAergic and nicotinic signaling.

### GABAergic control over DA axon excitability

Our results are in agreement with previous studies that have reported a striatal GABAergic tone (Gruen et al., 1992; Ade et al., 2008; Koh et al., 2023; Day et al., 2024). The striatum is a predominantly GABAergic structure consisting of mostly GABAergic medium spiny neurons along with a variety of GABAergic interneurons, each with a unique connectivity pattern to the surrounding circuitry (Taverna et al., 2008; Dobbs et al., 2016; Tepper et al., 2018; Dorst et al., 2020; Holly et al., 2021; Kocaturk et al., 2022). As a result, there are many possible sources of GABA within the striatum, including these GABAergic neurons, non-neuronal cells, co-release from DA axons, and co-release from CINs – any combination of which could be contributing to the effects observed here (Tritsch et al., 2012; Wójtowicz et al., 2013; Lozovaya et al., 2018; Roberts et al., 2020; Patel et al., 2024). Recently, Patel and colleagues showed that GABA co-released from DA axons can inhibit the same axons via autoregulatory GABA_A_ receptors (Patel et al., 2024). Together with the data presented here and past results showing that CINs can drive GABA release from DA axons (Nelson et al., 2014), this feature could endow the CIN-DA axon striatal circuit with a complex means of intrinsic negative feedback.

The results presented here are also consistent with GABA_A_ receptors exerting a shunting effect on DA axons, as we described previously (Kramer et al., 2020). Shunting inhibition is defined as a decrease in both input resistance and as a result, membrane space constant, both of which would predict the slower and smaller evoked axEPSPs observed in diazepam (Farrant & Kaila, 2007). Both functional and morphological data suggest that DA axons likely receive nicotinic input from release sites located throughout the axonal arbor, and a specific site for ectopic AP initiation has not been identified (Chang, 1988; Jones et al., 2001; C. Liu et al., 2022; Kramer et al., 2022). This means that subthreshold axEPSPs are likely integrated both across time (i.e. following synchronous CIN firing) and space (i.e. following activation of different nAChRs spread along the length of the DA axon) in order to summate and evoke APs and therefore could be especially prone to regulation by GABA_A_ receptor-mediated shunting inhibition.

Overall, this work offers a possible explanation for the different effects of diazepam on DA release with and without nAChR antagonists that we reported previously (Kramer et al., 2020). With nicotinic transmission intact, the majority of electrically-evoked DA release and evoked presynaptic Ca^2+^ in DA axons arises from nAChR activation and subsequent APs (Threlfell et al., 2012; Cachope et al., 2012; C. Liu et al., 2022). Our previous results explain the moderate effects of diazepam in the pharmacologically isolated condition and highlight the direct influence of GABA_A_ receptors on DA axon excitability (Kramer et al., 2020). Our new findings suggest that GABA_A_ receptors additionally regulate DA axon excitability via control of nicotinic input, as illustrated by the decreased amplitude and slowed kinetics of axEPSPs in diazepam. These data suggest that diazepam will also decrease nAChR-evoked DA release, leading to the larger effect observed with nicotinic signaling intact (Kramer et al., 2020).

Importantly, although GABA_A_ receptors on CINs could be exerting additional effects on the circuitry, our results suggest that electrically-evoked ACh release is not influenced by GABA_A_ receptor activation. Altogether, these data fit with the hypothesis that axEPSPs are shunted by axonal GABA_A_ receptor activation leading to impaired integration in DA axons. Our collection of evidence suggests that GABA_A_ receptor-mediated inhibition of DA release occurs both due to changes in intrinsic excitability of DA axons and due to the diminished effect of nicotinic input, suggesting an even more robust form of control than previously described (Lopes et al., 2019; Kramer et al., 2020).

### Dopaminergic axons as integrators of striatal information

A growing body of literature has described a variety of presynaptic receptors as poised to modulate DA release from the densely arborized and unmyelinated striatal DA axons (Matsuda et al., 2009; Threlfell et al., 2012; Cachope & Cheer, 2014; Aransay et al., 2015; Kramer et al., 2020, 2022). While it is well known that DA axons relay somatodendritic information via propagated APs, these studies have led to accumulating evidence for an additional role of DA axons through local, modulatory control of DA release (Rice & Cragg, 2004; H. Zhang & Sulzer, 2004; Threlfell et al., 2012; C. Liu et al., 2022; Kramer et al., 2022). Specifically, the recent description of spontaneous nicotinic axEPSPs and nAChR-evoked APs has provided a mechanistic explanation for how these receptors are able to shape axonal physiology in *ex vivo* slices (C. Liu et al., 2022; Kramer et al., 2022). These findings suggest the possibility that DA axons may function in ways canonically thought to be unique to dendrites, such as converting subthreshold excitatory input into APs. Our results provide evidence for yet another function of DA axons that is reminiscent of the somatodendritic region: integration of inhibition and excitation to shape output.

Although the biophysical factors controlling local axonal AP initiation are not specifically known, conditions that favor fast, synchronous, large-amplitude depolarizations will presumably increase the likelihood of subthreshold input summating into APs. This points to the intriguing possibility that GABA_A_ receptor activation could control how much influence CINs have over local DA release. Specifically, the local GABAergic tone could dictate the extent to which CIN synchrony is necessary to evoke APs in DA axons. Additionally, a striatal GABAergic tone could compartmentalize the widely arborized axons, allowing for potentially more complex computations as is observed with dendritic compartmentalization (Trigo et al., 2008; Wybo et al., 2019). Importantly, changes to tonic GABAergic inhibition in the striatum have been reported in mouse models of neurodegenerative diseases, namely Parkinson’s Disease (PD) and Huntington’s Disease (Cepeda et al., 2013; Wójtowicz et al., 2013; Roberts et al., 2020). For example, Roberts and colleagues recently showed that in the absence of cholinergic influence, tonic GABAergic inhibition of DA axons is substantially increased exclusively in the dorsal, but not ventral, striatum of an early-disease model of PD (Roberts et al., 2020). The interplay between these disease-related changes and well-documented changes to CINs in PD could be an interesting avenue of investigation, with potential therapeutic value (DeBoer et al., 1993; Ding et al., 2006; Pisani et al., 2007; McGregor & Nelson, 2019; Nielsen & Ford, 2023).

Here, we provide evidence that DA axons in the striatum integrate multiple inputs from their local environment. Specifically, the combination of tonic extracellular GABA and phasic ACh input contributes to the output of DA axons via the regulation of local initiation of APs. Our data show that the effects of GABA_A_ receptor activation on nicotinic input to DA axons are not due to circuit mechanisms. Instead, we propose that DA axons may perform integration-like computations – a feature that has not previously been observed in axonal structures but is characteristic of postsynaptic dendrites. Additionally, we show that picrotoxin exerts robust off-target effects on nAChRs and should be used carefully in the striatum and other complex circuitry. Finally, we describe a photometry analysis method that combines genetic specificity with high temporal resolution in order to study presynaptic excitability. Overall, our data provide evidence for even more nuanced local control of DA axon physiology than previously understood, and we suggest a novel approach to study this phenomenon and similar ones across brain regions.

## METHODS

### MICE

All animal handling and procedures were approved by the Animal Care and Use Committee (ACUC) for the National Institute of Neurological Disorders and Stroke (NINDS) at the National Institutes of Health. All experiments were performed on adult mice between the ages of p51 and p143 and mice of both sexes were used throughout the study. For patch clamp experiments, DAT-Cre (RRID:IMSR_JAX:006660) mice were crossed with Ai9 (RRID:IMSR_JAX:007909) mice and the resulting DAT-Cre::Ai9 offspring were used. For imaging experiments, DAT-Cre mice underwent stereotaxic injections at postnatal day 39 or older and were used between 2 and 11 weeks later.

### VIRAL INJECTIONS

For stereotaxic injections, mice were maintained under isoflurane for the duration of the procedure. The viruses used for this study were jGCaMP8s (AAV9-pGP-syn-FLEX-jGCaMP8s-WPRE titer: > 1x 10^13^, Addgene #162377) and GRAB_ACh3.0_ (AAV9-hSyn-DIO-GRABACh3.0 titer: > 1x 10^12^, plasmid: Addgene #121923, virus packaged by Vigene). Viral aliquots were injected (0.1-0.5 μL) bilaterally into the SNc (X: ± 1.9 Y: -0.5 Z: -3.9) via a Hamilton syringe. At the end of the injection, the needle was raised at a rate of 0.1 to 0.2 mm per minute for 10 minutes before being removed. Mice were allowed to recover from anesthesia on a warmed pad and returned to the home cage.

### TISSUE SLICING

Mice were anesthetized with isoflurane, decapitated, and brains were rapidly extracted. Horizontal sections were cut at 320-380 μm thickness on a vibratome while immersed in warmed (34°C), modified, slicing ACSF containing (in mM) 198 glycerol, 2.5 KCl, 1.2 NaH_2_PO_4_, 20 HEPES, 25 NaHCO_3_, 10 glucose, 10 MgCl_2_, 0.5 CaCl_2_. Cut sections were promptly removed from the slicing chamber and incubated for 30-60 minutes in a heated (34°C) chamber with holding solution containing (in mM) 92 NaCl, 30 NaHCO_3_, 1.2 NaH_2_PO_4_, 2.5 KCl, 35 glucose, 20 HEPES, 2 MgCl_2_, 2 CaCl_2_, 5 Na-ascorbate, 3 Na-pyruvate, and 2 thiourea. Slices were then stored at room temperature and used 30 min to 6 hours after slicing. Following incubation, slices were moved to a heated (33–35°C) recording chamber that was continuously perfused with recording ACSF (in mM): 125 NaCl, 25 NaHCO_3_, 1.25 NaH_2_PO_4_, 3.5 KCl, 10 glucose, 1 MgCl_2_, 2 CaCl_2_.

### ELECTROPHYSIOLOGY AND FLUORESCENCE IMAGING

For imaging experiments, a white light LED (Thorlabs; SOLIS-3C) was used in combination with a EGFP (Chroma; 49002) filter set to visualize DA axons infected with jGCaMP8s or GRAB_ACh3.0_. To visualize jGCaMP8s or GRAB_ACh3.0_ signals, a photodiode (New Focus) was mounted on the top port of the Olympus BX-51WI. Signals were acquired every 30 seconds (jGCaMP8s) or 60 seconds (GRAB_Ach_) using a Digidata 1440A (Molecular Devices) sampled at 50kHz. All electrical stimulation was delivered with tungsten bipolar electrodes (250 μm tip separation, MicroProbes) placed 200 μm from the imaging site in the DMS. Electrical stimulation was delivered using an Isoflex (A.M.P.I.) with amplitudes ranging from 0.3 to 0.6 V to produce a near-maximal (∼90-95% of maximum) signal for photometry experiments, and a subthreshold signal for perforated patch experiments.

Perforated-patch recordings from striatal TdTomato+ axons were made using borosilicate pipettes (6-9 MΩ) filled with internal solution containing (in mM) 135 KCl, 10 NaCl, 2 MgCl_2_, 10 HEPES, 0.5 EGTA, 0.1 CaCl_2_, adjusted to a pH value of 7.43 with KOH, 278 mOsm. Pipette tips were backfilled with ∼1 µL of gramicidin-free internal. Pipettes were then filled with internal containing between 80 and 100 µg/mL gramicidin. Patch integrity was monitored by the addition of Alexa-488 to the gramicidin-containing internal. Whole-cell recordings from the TdTomato+ axons in the medial forebrain bundle were made using borosilicate pipettes (4-7 MΩ) filled with internal solution containing (in mM) 122 KMeSO_3_, 9 NaCl, 1.8 MgCl_2_, 4 Mg-ATP, 0.3 Na-GTP, 14 phosphocreatine, 9 HEPES, 0.45 EGTA, 0.09 CaCl_2_, adjusted to a pH value of 7.35 with KOH. All recordings were made with a MultiClamp 700B (Molecular Devices).

Pressure ejection of acetylcholine was performed using borosilicate pipettes (2-4 MΩ). Acetylcholine (300 µM) was added to a modified external solution containing (in mM): 125 NaCl, 25 NaHCO_3_, 1.25 NaH_2_PO_4_, 3.5 KCl, 10 HEPES, 0.01 Alexa 488, final osmolarity 280–290 mOsm. This puffing solution was then spin filtered, loaded into a glass pipette, and lowered to within 30–50 µm of the axon using a micro-manipulator. The puffing solution was applied onto the axon with a short pressure ejection (100–250 msec in duration) using a PV 820 Pneumatic PicoPump (WPI). In a subset of puffing experiments, TTX (500 nM) was included in the bath in order to eliminate APs propagating from the soma. In other cases, propagating APs were removed *post hoc* to aid in analysis of puff-evoked events.

### QUANTIFICATION AND STATISTICAL ANALYSIS

Analysis was conducted in Igor Pro (Wavemetrics) and statistical tests were performed in Prism 9 (GraphPad) and Igor Pro. Data in text are reported as the mean +/- the standard error of the mean. T-tests were used for two-group comparisons, and ANOVA tests were used when comparing more than two groups followed by a Bonferroni post-hoc test for analysis of multiple comparisons.

#### axEPSP analysis

For amplitude measurements, evoked axEPSPs from every trial recorded were measured relative to baseline prior to the electrical stimulus. The data from each axon was normalized to its average amplitude during the control period. To perform measurements of axEPSP slope, average waveforms were generated for each axon in both control conditions and following application of diazepam. The rising slope of each average waveform was calculated from 30% to 70% of peak amplitude, and the slope of the diazepam waveform was normalized to that of the control waveform for each axon recorded.

#### Discrimination between direct and nicotinic components

We determined that the axonal Ca^2+^ signal was comprised on two components: a component resulting from the direct stimulation of dopaminergic axons (“direct component”) and a delayed component that was dependent upon the activation of axonal nAChRs (“nicotinic component”). For discrimination between the direct and nicotinic components, analysis windows were defined based on average waveforms. Specifically, the direct component window was defined as a region ranging from 1.1-6.7 msec following the electrical stimulus. The window for the nicotinic component analysis was defined as a 20 msec window immediately following the direct component (i.e. 6.7-36.7 msec after electrical stimulation; Fig S1).

#### Amplitude quantification of direct and nicotinic components

In order to quantify changes to the direct component of evoked jGCaMP8s signals, the first derivative of the raw signal was lightly smoothed. The maximum value within the direct component analysis window was determined for every trace in each slice to create a time series. To quantify the nicotinic component, we determined the DHβE-sensitive component by subtracting the average waveform in DHβE from each control trace. To create the time series, the peak amplitude of each trace was measured from baseline and adjusted to account for baseline noise. Finally, both the quantifications of direct and adjusted nicotinic components for each slice were normalized to control conditions and averaged across slices. Quantification of drug effects was done by averaging the final 5 minutes of the control or wash-in periods.

#### Quantification of direct and nicotinic component kinetics

To find the onset latency of the direct component, the differentiated traces were used. Onset was defined for each trace as the point at which the trace crossed 10% of the maximum amplitude of the direct component. This point was identified by a backwards search beginning at the end of the direct component analysis window. The latency to peak of the nicotinic component was found by using the nicotinic component traces described above (Fig. S2).

## ACKNOWLEDGMENTS

We thank Drs. Anna Lipkin, Jessica Perkins, Lorenzo Sansalone, and Dana Cobb-Lewis of the Khaliq Laboratory for their insightful discussions and comments on this manuscript. This work was supported by NINDS Intramural Research Program grant NS003135 to Z.M.K. Funding for this work was also provided by the Aligning Science Across Parkinson’s (ASAP) Collaborative Research Network through a grant to Z.M.K.

## AUTHOR CONTRIBUTIONS

Conceptualization: S.G.B.-W., P.F.K., and Z.M.K.; Methodology: S.G.B.-W., P.F.K., and Z.M.K.; Investigation: S.G.B.-W., P.F.K., A.Y., F.H.C., and R.Z.; Validation: S.G.B.-W., P.F.K., and Z.M.K.; Software: S.G.B.-W. and P.F.K.; Formal analysis: S.G.B.-W. and Z.M.K.; Data curation, S.G.B.-W., P.F.K., and Z.M.K.; Visualization, S.G.B.-W. and Z.M.K.; Resources, supervision, project administration, and funding acquisition: Z.M.K.; Writing – original draft: S.G.B.-W.; Writing – reviewing and editing: S.G.B.-W., P.F.K., and Z.M.K.

---

**Supplemental Figure 1:**
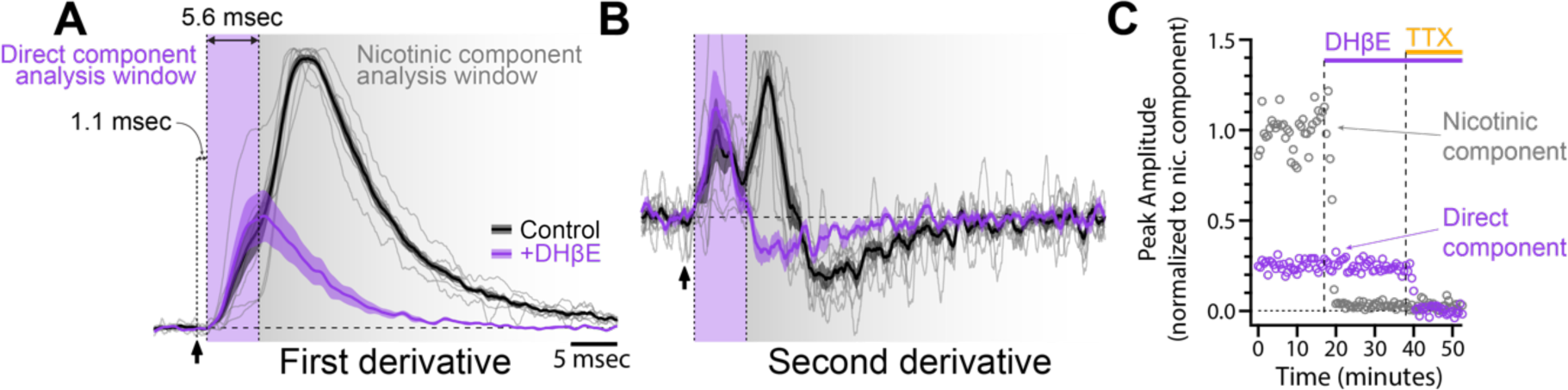
Definition of direct and nicotinic components in jGCaMP8s photometry traces. **(A)** First derivative of evoked jGCaMP8s signals normalized to the peak during control trials. Thin lines represent data from 8 individual experiments. The black trace is the average of all experiments during the control condition, and the purple trace is the average of the same experiments following application of DHβE (1 μM). Analysis windows are defined based on traces in (B). **(B)** Second derivative of the traces shown in (A). Direct component analysis window (purple region) begins at the onset of the first peak and ends at the onset of the second peak, when the nicotinic component analysis window (grey region) begins. **(C)** Time course of the direct and nicotinic component for the slice shown in Fig. 2B and E during DHβE and TTX wash-in.

**Supplemental Figure 2:**
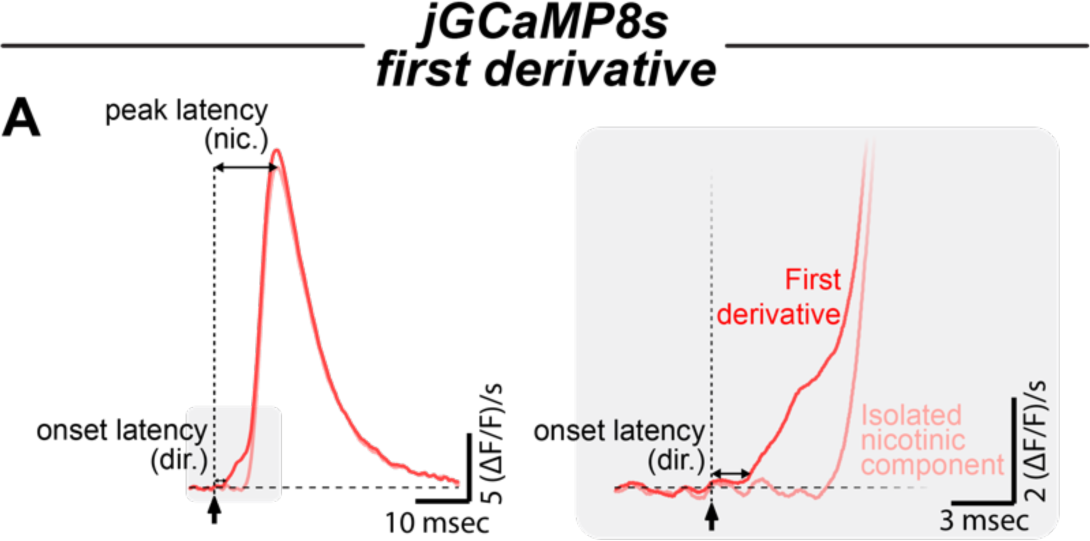
Explanation of latency measurements for direct and nicotinic components. **(A)** Illustration of where measurements were made for Fig. 3G. Peak latency of the nicotinic component was defined as the time at which the subtracted trace reached its maximum value. Onset latency of the direct component was defined as the time at which the first derivative trace crossed 10% of the direct component peak amplitude. Onset latency was used for the direct component because the peak of the component did not always fall within the boundaries of the analysis window.

**Supplemental Figure 3:**
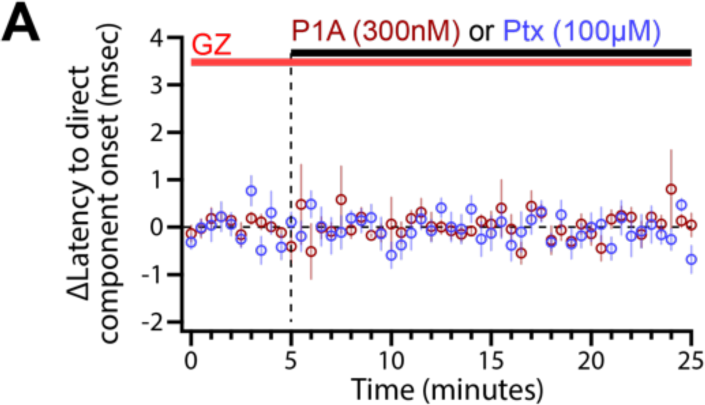
**Direct component onset latency is not impacted by P1A or picrotoxin**. **(A)** Time course of the onset latency of the direct component following application of either picrotoxin (n = 8) or P1A (n = 5). All slices were bathed in ACSF containing 10 μM gabazine for the duration of the experiment. From the same experiments as Fig. 6G.

